# Heterogeneity of *Candida* bloodstream isolates in an academic medical center and affiliated hospitals

**DOI:** 10.1101/2025.02.05.636768

**Authors:** Nancy E. Scott, Elizabeth Wash, Christopher Zajac, Serin E. Erayil, Susan E. Kline, Anna Selmecki

**Affiliations:** University of Minnesota, Bioinformatics and Computational Biology Program; University of Minnesota, Department of Microbiology and Immunology; University of Minnesota, Molecular, Cellular, Developmental Biology and Genetics Program; University of Minnesota, Department of Medicine, Division of Infectious Diseases and International Medicine

**Keywords:** Candidemia, antifungal drug resistance, emerging *Candida* species, serial isolates

## Abstract

Invasive *Candida* bloodstream infections (candidemia) are a deadly global health threat. Rare *Candida* species are increasingly important causes of candidemia and phenotypic data, including patterns of antifungal drug resistance, is limited. There is geographic variation in the distribution of *Candida* species and frequency of antifungal drug resistance, which means that collecting and reporting regional data can have significant clinical value. Here, we report the first survey of species distribution, frequency of antifungal drug resistance, and phenotypic variability of *Candida* bloodstream isolates from an academic medical center and 5 affiliated hospitals in the Minneapolis-Saint Paul region of Minnesota, collected during an 18-month period from 2019 to 2021. We collected 288 isolates spanning 11 species from 119 patients. *C. albicans* was the most frequently recovered species, followed by *C. glabrata* and *C. parapsilosis*, with 10% of cases representing additional, rare species. We performed antifungal drug susceptibility for the three major drug classes and, concerningly, we identified fluconazole, micafungin and multidrug resistance rates in *C. glabrata* that were ∼ 2 times higher than that reported in other regions of the United States. We report some of the first phenotypic data in rare non-*albicans Candida* species. Through analysis of serial isolates from individual patients, we identified clinically relevant within-patient differences of MIC values in multiple drug classes. Our results provide valuable clinical data relevant to antifungal stewardship efforts and highlight important areas of future research, including within-patient dynamics of infection and the mechanisms of drug resistance in rare *Candida* species.

## INTRODUCTION

*Candida* species are frequent human commensals and also important opportunistic fungal pathogens^1–3^. *Candida* infections can be superficial, such as oral candidiasis, or deeply invasive, including sites like the bloodstream (candidemia) or abdominal cavity. About 1.5 million cases of invasive candidiasis occur annually around the world^4^.

Diverse *Candida* species are increasingly important causes of invasive candidiasis, and while *Candida albicans* is the most frequent cause, its global prevalence has decreased to less than 50% of reported cases^5^. *Candida* species continue to undergo nomenclature changes; therefore we will use the species names most familiar to clinicians, with the revised name in parentheses when the species is introduced. The most common non-*albicans Candida* pathogens include *C. glabrata* (*Nakaseomyces glabratus*), *C. parapsilosis* and *C. tropicalis*; prevalence of each species varies between global regions^5^. *C. glabrat*a is the second most common cause of invasive candidiasis in North America, Europe and Australia^5,6^. In Latin America *C. parapsilosis* follows *C. albicans* in frequency of isolation, except in Columbia and Venezuela, where *C. parapsilosis* is the most common cause of candidemia^7^. *C. tropicalis* accounts for ∼7.5% of invasive candidiasis in Europe and ∼17% of cases in Latin America^5,7^. A growing percentage of invasive infections are caused by rare species and emerging pathogens^8,9^. For example, *C. krusei* (*Pichia kudriavzevii*) is responsible for 2-3% of invasive candidiasis cases^8^. *C. auris* is a recently emerged pathogen that has spread globally and can be transmitted between patients^10^. *C. lusitaniae* (*Clavispora lusitaniae*) is closely related to *C. auris* and accounts for 2-3% of invasive candidiasis cases^9,11^.

Major antifungal drug classes are limited to azoles, echinocandins and polyenes. Azoles such as fluconazole, voriconazole and itraconazole cause cell membrane stress and are fungistatic^12^. Azoles target Erg11, part of the ergosterol biosynthesis pathway^12^. Echinocandins such as micafungin, caspofungin and anidulafungin, are fungicidal and target the Fks subunit of 1,3-beta-D-glucan synthase which results in cell wall stress^13,14^. The fungicidal polyenes, including amphotericin B, target ergosterol in the cell membrane leading to cell membrane stress^15^. Some *Candida* species have intrinsic resistance to specific antifungal drugs – for example, *C. krusei*’s Erg11 protein has naturally reduced susceptibility to fluconazole^16,17^. *Candida* species also acquire drug resistance through a broad spectrum of mutations. Mechanisms of azole resistance are diverse and include overexpression or mutation of the drug target Erg11p, alteration of the ergosterol pathway and increased activity of drug efflux pumps^18–20^. Echinocandin resistance is driven primarily by mutations in the gene(s) encoding the Fks subunit of the drug target^21^.

The frequency of acquired antifungal drug resistance varies between species, drug class and geographic regions^5,7,9^. Fluconazole resistance is more frequent in non-*albicans Candida* species, including *C. glabrata* (∼9%), *C. tropicalis* (∼9 – 12%) and *C. auris* (∼90%)^5,22^. Echinocandin resistance in *C. glabrata* is higher in North America (2.8%), compared to Europe (0.6%) and the Asia-Pacific region (0.4%)^5^. Resistance rates can also vary by institution, e.g. echinocandin resistance rates of *C. glabrata* isolates range from 0 to 25% within different hospitals in the United States^23^. Multidrug resistance (MDR), defined as resistance to more than one class of antifungal drug, is a growing concern. *C. auris* is best known for rapid acquisition of multidrug resistance, but acquired multidrug resistance has also been reported in *C. albicans*, *C. glabrata*, *C. parapsilosis*, *C. tropicalis*, *C. krusei*, and *C. lusitaniae*^24–29^. With only limited options available to treat invasive *Candida* infections, it is critical to understand the species distribution and frequency of antifungal drug resistance at a local level to make appropriate therapeutic choices and prevent future outbreaks.

Rare *Candida* species lack sufficient clinical data to accurately define antifungal susceptibility cutoff values. Susceptibility testing of an isolate identifies the minimum inhibitory concentration (MIC) value in a given drug. Setting a clinical breakpoint (i.e., the MIC value at which isolates are considered resistant) requires testing large numbers of isolates to determine the distribution of MIC values for a given species and then integrating MIC results with clinical outcome data^30^. As a result, clinical breakpoints so far are limited to common pathogens such as *C. albicans* or *C. glabrata*^31^. In the absence of clinical breakpoints, epidemiological cut-off values (ECOFFs) have been determined for some additional species^32^. ECOFF values define the upper limit of the ‘wild-type’ MIC distribution for a species and are based on MIC testing of multiple, independent groups of isolates^32,33^. ECOFFs can provide information about the expected drug response of a species when breakpoints have not been established due to insufficient clinical evidence (i.e., limited treatment outcome data). Gathering more data from rare and emerging *Candida* pathogens is crucial to develop ECOFFs and clinical breakpoints which can guide treatment decisions.

In the absence of antifungal drug resistance, some isolates demonstrate drug tolerance: persistent growth in drug concentrations above their MIC^34,35^. Tolerance is distinct from resistance, and increased fluconazole tolerance in *C. albicans* has been associated with failure to clear an infection during extended therapy^35^. For fungistatic drugs such as fluconazole, one measure of tolerance is supra-MIC growth (SMG)^36^. Tolerance has been most studied in *C. albicans*, however the extent of azole tolerance across clinical isolates is poorly understood. Tolerance levels in non-*albicans Candida* species is not known and might impact patient outcomes across species^37,38^.

Serial clinical isolates from an individual patient can display phenotypic variation including changes in antifungal drug resistance^39^. Most previous studies of clinical *Candida* strains focused on only one isolate per patient^40–42^. Few studies have analyzed serial clinical isolates and the extent and impact of within-host variation of *Candida* populations on clinical outcomes is poorly understood^39,40,43^.

Surveys of invasive candidiasis and candidemia that only examine data at the level of continent or country do not account for important regional variation^44–46^. For example, in 2016, the CDC’s Emerging Infections Program (EIP) reported that the prevalence of *C. albicans* in candidemia cases ranges from 35% in Maryland to 42% in Tennessee, while the prevalence of *C. parapsilosis* ranges from 9% in Tennessee to 18% in Oregon^9^. The EIP’s 2012 – 2016 candidemia surveillance only included metro areas from four states – Georgia, Maryland, Oregon and Tennessee^9,45^. To our knowledge, no studies have reported the species distribution of candidemia cases in Minnesota’s Twin Cities region.

To investigate the species distribution, frequency of antifungal drug resistance, and phenotypic variability of Minneapolis – St. Paul (Twin Cities) metro area *Candida* bloodstream infections, we prospectively collected residual clinical bloodstream isolates from an academic medical center and 5 affiliated hospitals in the Twin Cities metro area during an 18-month period from 2019 to 2021. Our isolate bank includes a total of 288 isolates from 119 patients, with 11 *Candida* species represented. We performed antifungal susceptibility testing of all isolates for each of the three major drug classes. We identified multidrug resistance in a single *C. lusitaniae* isolate, and troublingly, in 4.8% of *C. glabrata* isolates. By collecting serial isolates from individual patients, we identified within-host differences in MIC values and the acquisition of multidrug resistance. Additionally, we provide some of the first data related to tolerance in non-*albicans Candida* species. Our study is the first to report the diversity of candidemia-causing species and frequency of antifungal drug resistance in a major hospital system in the Twin Cities metro area of Minnesota.

## RESULTS

### *Candida albicans* is the most common cause of candidemia in the Twin Cities area

We collected isolates from all positive *Candida* blood cultures identified during clinical testing between December 2019 and May 2021 (see Methods and Table S1). For this study, we define an isolate as a single colony subculture taken from an individual blood culture sample (Figure 1A). We collected a total of 288 isolates representing 11 species from 119 different patients (Figure 1B and C). *C. albicans* was the most frequently identified species in the study and was isolated from 54 patients (45.3%), followed by *C. glabrata* (n=42, 35.3%), *C. parapsilosis* (n = 8, 6.7%) and *C. tropicalis* (n = 3, 2.5%). Rare species detected in this study include *C. dubliniensis*, *C. kefyr* (*Kluyveromyces marxianus*), *C. orthopsilosis*, and *C. lusitaniae* (each isolated from 3 patients), *C. krusei* (2 patients), *C. nivariensis* (*Nakaseomyces nivariensis*, 1 patient) and *C. utilis* (*Cyberlindnera jadinii*, 1 patient).

**Figure 1.**
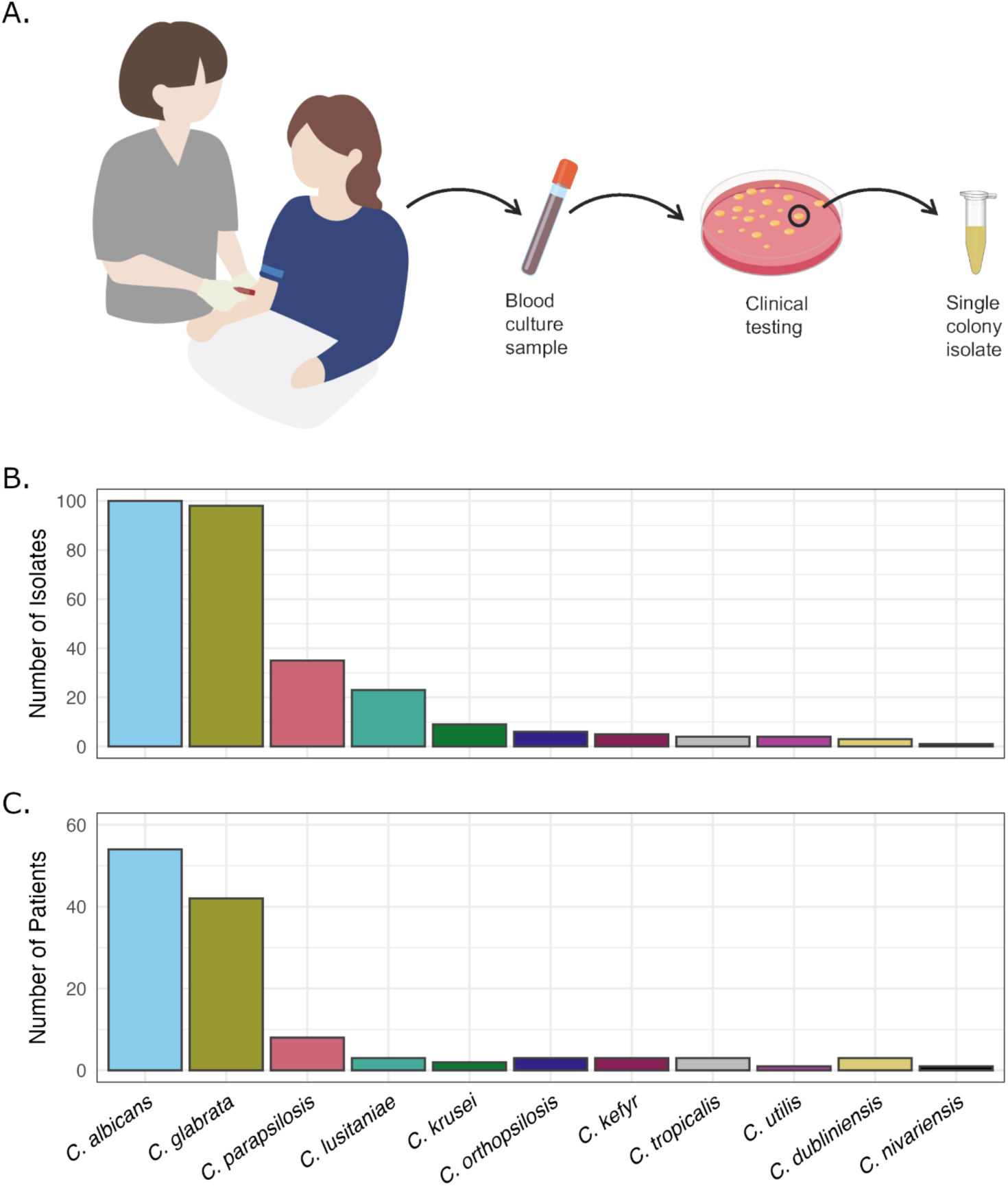
Candidemia isolate collection summary. (A). Workflow for isolate collection. Positive blood culture samples were subcultured for microbial identification as part of the clinical workflow. Species-level identification was performed by matrix-assisted laser desorption/ionization time-of-flight (MALDI-TOF). *Candida* species were flagged by clinical staff and a single colony was selected from the initial plate to be cultured and saved for our study. (B) Number of isolates collected per *Candida* species. (C) Number of patients with a bloodstream infection of each *Candida* species.

### Multiple isolates from individual patients demonstrate the within-host diversity of clinical strains

Forty-eight percent of patients in our study had multiple positive blood cultures during the study period and we collected one isolate from each positive culture. We defined a case as all isolates collected from an individual patient, and each case was assigned a numeric code that was unrelated to any patient identifiers. To provide more detailed information about isolates collected throughout individual patient infections, we further defined four categories of cases: 1) individual cases (one isolate collected from one patient); 2) Serial isolate cases (multiple isolates of a single species collected from one patient within 30 days of the initial positive culture); 3) Recurrent cases (multiple isolates of a single species, collected from one patient more than 30 days after the initial positive culture); and 4) Polyfungal cases (multiple species collected from a single patient within 30 days of each other). Serial isolate cases were collected from all species except *C. dubliniensis* and *C. nivariensis* (Figure 2 and Table S2). Four recurrent cases were identified (3.4% of all cases); two were recurrent *C. albicans* infections and two were recurrent *C. parapsilosis* infections. The time span between sampling of recurrent isolates ranges from 107 to 338 days (Figure 2, Table S3).

**Figure 2.**
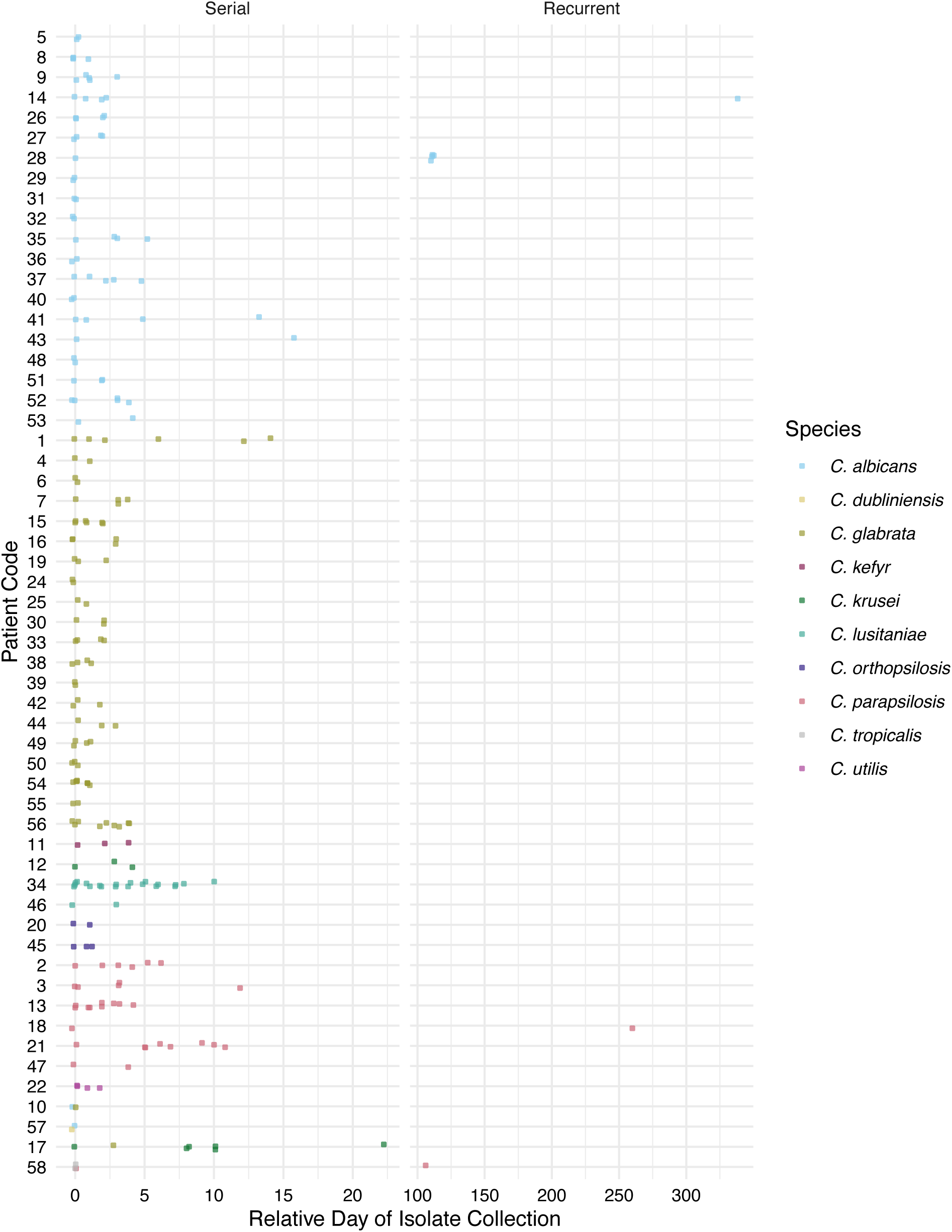
Timelines of serial, recurrent and polyfungal isolate sampling. Patient codes are on the y-axis and the relative number of days are on the x-axis. Serial cases have multiple isolates of a single species collected from one patient within 30 days of the initial positive culture. Recurrent cases have additional isolates of the same species collected more than a month after the initial positive blood culture.

We identified four polyfungal cases (Figure 2, Table S3), including two cases that each involved two different species collected independently on one day (*C. albicans* and *C. glabrata* isolated from patient 10; *C. albicans* and *C. dubliniensis* isolated from patient 57). We found no patterns related to the species which were isolated together or the time spans involved in polyfungal cases. Two of our patients fit into multiple categories, highlighting the complexity of some candidemia infections. Patient 17 had six *C. krusei* blood cultures collected over 22 days, with an additional single *C. glabrata* blood culture on the third day, comprising both a serial and polyfungal case. Patient 58 had two independent *C. tropicalis* blood cultures and a *C. parapsilosis* blood culture collected over the course of two days, and another *C. parapsilosis* blood culture over three months later, therefore fitting the categories of serial, polyfungal and recurrent cases.

### Antifungal resistance is most common in *C. glabrata* but also occurs in other non-*albicans* species

We performed antifungal susceptibility testing on all isolates to determine the frequency of resistance against the three major antifungal drug classes. We measured the minimum inhibitory concentration (MIC) for fluconazole, micafungin and amphotericin B using the European Committee on Antimicrobial Susceptibility Testing (EUCAST) broth microdilution method and interpreted results according to EUCAST breakpoints (Supplementary Table S1)^31,33^. For 8 of 288 isolates that had insufficient growth for EUCAST growth criteria, we measured the MIC by gradient diffusion (Methods). For species without clinical resistance breakpoints, we evaluated available ECOFF values.

Fluconazole clinical resistance breakpoints were available for all species in this study except for *C. krusei*, which has intrinsic resistance. We identified 22 *C. glabrata* isolates, 1 *C. tropicalis* isolate and 4 *C. utilis* isolates with fluconazole resistance (Figure 3, Supplementary Table S1). The clinical breakpoint of fluconazole for *C. glabrata* is 16 μg/mL, and MIC values for resistant *C. glabrata* isolates ranged from 32 to 256 μg/mL. All *C. albicans*, *C. parapsilosis*, *C. lusitaniae*, *C. orthopsilosis*, *C. kefyr*, *C. dubliniensis* and *C. nivariensis* isolates were clinically susceptible to fluconazole.

**Figure 3.**
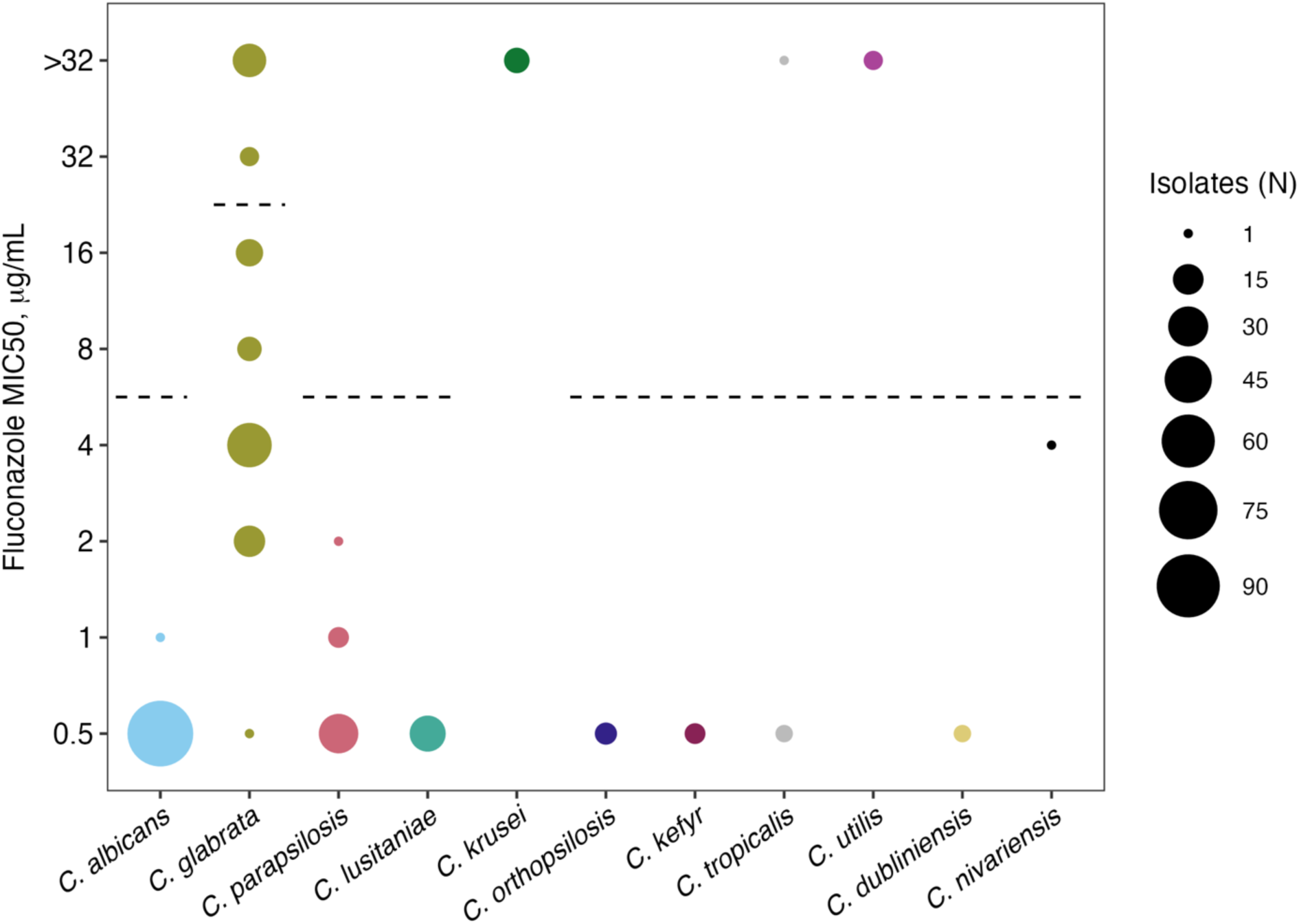
Summary of fluconazole MIC values for all isolates. The size of each circle represents the number of isolates with that MIC. Established clinical resistance breakpoints are indicated by horizontal dashed lines. 33 *C. glabrata* isolates, 1 *C. tropicalis*, and 4 *C. utilis* isolates are fluconazole resistant. Fluconazole MIC screening was performed up to a maximum concentration of 32 μg/mL, which exceeds clinical breakpoints, and is shown in this figure. *C. glabrata*, *C. tropicalis* and *C. utilis* isolates with fluconazole MIC values > 32 μg/mL were subsequently tested at higher concentrations to determine MIC values which are reported in the text and Table S1.

Micafungin clinical resistance breakpoints are only established for *C. albicans*, *C. glabrata*, and *C. parapsilosis*. We identified 11 micafungin resistant *C. glabrata* isolates (Figure 4, Table S1). All *C. albicans* have MIC values of <0.03 μg/mL and are micafungin sensitive. All *C. parapsilosis* isolates have MIC values of ≤ 2 μg/mL and are micafungin sensitive. All of our *C. krusei* and *C. tropicalis* isolates have MICs below their established ECOFF values (0.25 μg/mL for *C. krusei* and 0.06 μg/mL for *C. tropicalis)*, meaning that they fall within the wild-type MIC distribution^47^. No clinical breakpoints or ECOFF values are available for *C. lusitaniae, C. dubliniensis*, *C. kefyr*, *C. nivariensis*, *C. utilis* and *C. orthopsilosis*, but other surveys of clinical isolates have been reported. For example, *C. lusitaniae* clinical isolate MIC values ranging from 0.032 to 0.064 μg/mL have been reported previously^48,49^. Strikingly, 8 of our 23 *C. lusitaniae* isolates have MIC values ranging from 0.125 to > 1 μg/mL, indicating that they are micafungin resistant. Notably, we identified a range of resistant phenotypes within 1 serial case of *C*. *lusitaniae*, where 7 of 20 serial isolates from one patient had MIC values ranging from 0.256 to >1 μg/mL micafungin^50^. The micafungin MIC values for *C. dubliniensis*, *C. kefyr*, *C. nivariensis* and *C. utilis* in this study range from 0.016 to 0.064 µg/mL and are consistent with the median MIC values reported by other studies using EUCAST broth microdilution^49,51^. All *C. orthopsilosis* isolates have MIC values of 0.5 μg/mL, which is less than the median MIC values from other studies using EUCAST broth microdilution^51,52^.

**Figure 4.**
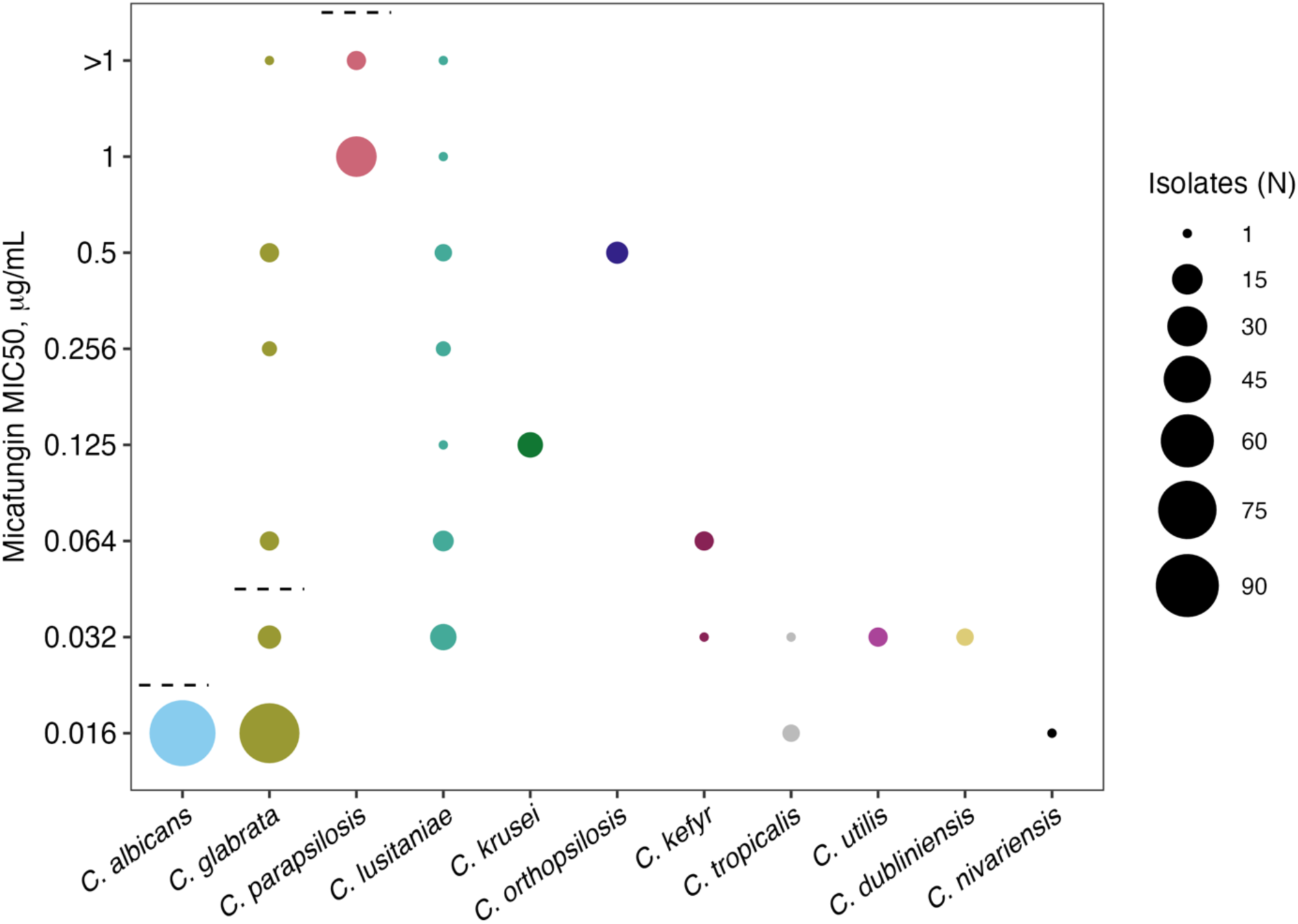
Summary of micafungin MIC values for all isolates. The size of each circle represents the number of isolates with that MIC. Established clinical resistance breakpoints are indicated by vertical dashed lines where available. 11 *C. glabrata* isolates and 8 *C. lusitaniae* isolates are micafungin resistant. Micafungin MIC screening was performed up to a maximum concentration of 1 μg/mL. *C. parapsilosis* isolates were subsequently tested at higher concentrations and all isolates were micafungin sensitive.

Amphotericin B clinical resistance breakpoints are available for *C. albicans*, *C. glabrata*, *C. parapsilosis*, *C. krusei*, *C. tropicalis* or *C. dubliniensis*. We did not find any amphotericin B resistance in these six species (Figure 5, Table S1). ECOFF values are available for *C. kefyr* (1 μg/mL) and *C. lusitaniae* (0.5 μg/mL)^53^. All *C. kefyr* isolates had MIC values below the ECOFF value. We identified a single amphotericin B-resistant *C. lusitaniae* isolate (MIC of 1 μg/mL). All *C. nivariensis, C. orthopsilosis* and *C. utilis* isolates tested for this study had MIC values of 1 μg/mL or less for amphotericin B. Since amphotericin B is reported to have similar *in vitro* activity among *Candida* species, this suggests that these isolates do not have any amphotericin B resistance^53^.

**Figure 5.**
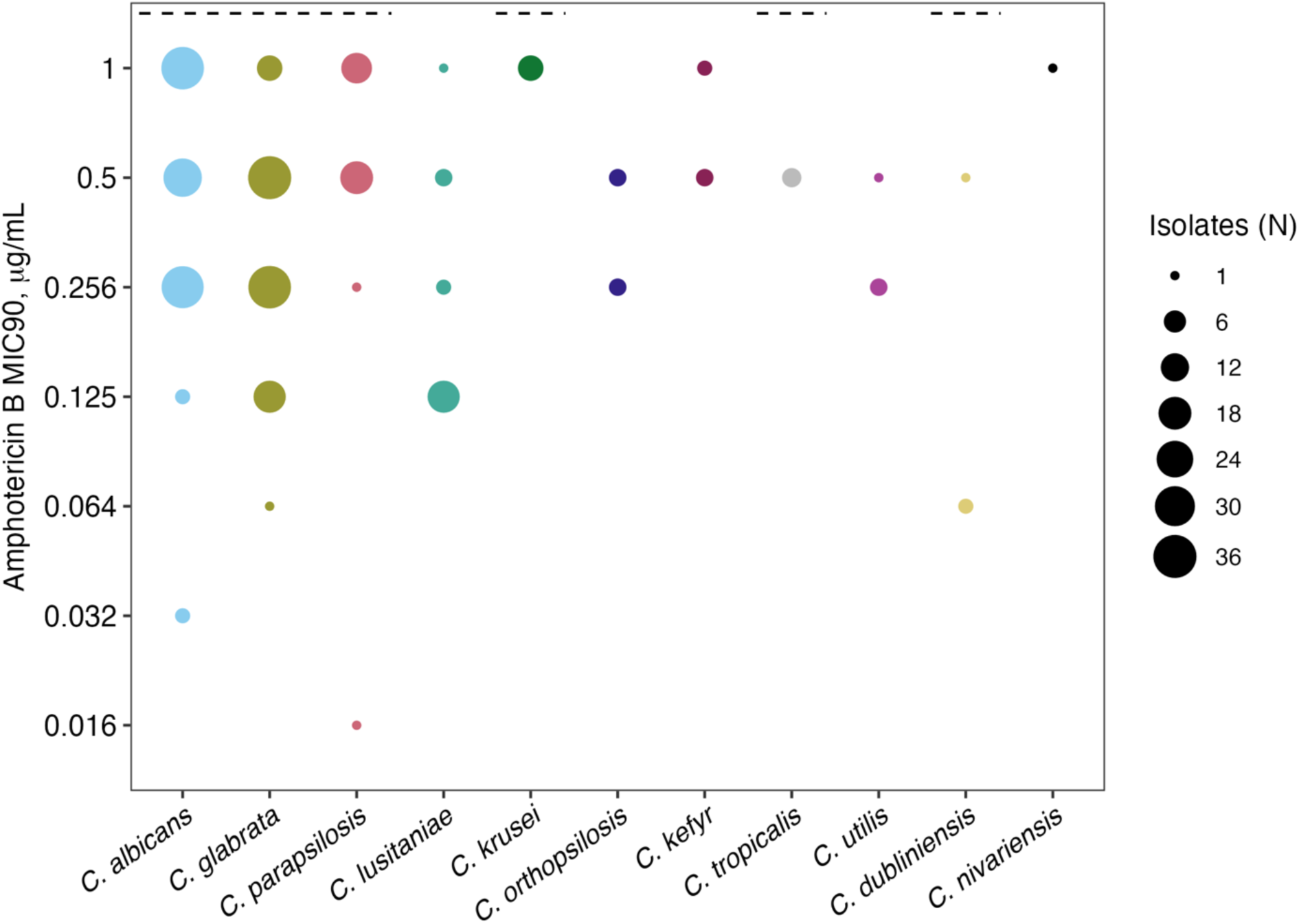
Summary of amphotericin B MIC values for all isolates. The size of each circle represents the number of isolates with that MIC. Established clinical resistance breakpoints are indicated by vertical dashed lines where available. One *C. lusitaniae* isolate is amphotericin B resistant.

### *C. glabrata* micafungin and fluconazole resistance are more frequent in our study compared to that reported from other regions of the United States

To determine the frequency of resistance at the case level, we determined the number of patients with any resistant isolates (i.e., serial resistant isolates from an individual patient count as a single case). The number and percentage of resistant cases per species is summarized in Table 1. Species with no resistant isolates are not listed.

**Table 1.** Frequency of antifungal drug-resistant cases by species.

| Species | Total cases, N | Drug | Resistant cases, N (%) |
| --- | --- | --- | --- |
| *C. glabrata* | 42 | fluconazole | 7 (16.7) |
|  |  | micafungin | 3 (7.1) |
| *C. lusitaniae* | 3 | micafungin | 2 (66.7) |
|  |  | amphotericin B | 1 (33.3) |
| *C. tropicalis* | 3 | fluconazole | 1 (33.3) |
| *C. utilis* | 1 | fluconazole | 1 (100) |

We found micafungin resistance in 7.1% of *C. glabrata* cases, about twice the frequency of cases resistant to any echinocandin (3.6%) reported by the CDC EIP for 2016^9^. We also identified fluconazole resistance in 16.7% of *C. glabrata* cases in our study, which is notably higher than the 10.7% reported by the CDC EIP for 2016^9^. We identified micafungin resistance in 2 of only 3 *C. lusitaniae* cases collected in our study. Overall, our results indicate that in *C. glabrata* antifungal drug resistance to two major drug classes is concerningly high in the Twin Cities metro area relative.

### Multidrug resistance is found in non-*albicans Candida* species

Multidrug resistance is an important clinical concern because antifungal treatment options are limited. We identified multidrug resistance in 33% of *C. lusitaniae* cases (n = 1 patient, micafungin and amphotericin B) and in 4.8% of *C. glabrata* cases (fluconazole and micafungin, n = 2 patients). The frequency of multidrug resistance in our study is almost two times the national average reported by the CDC EIP in 2016 (0 – 2.7%)^9^. Our results are concerning and important for informing local and national antifungal stewardship programs.

### Differences in MIC values between serial isolates occur in all three antifungal drugs

Clinical antifungal susceptibility testing is often only performed on the first isolate collected from a patient, limiting our understanding of both the within-host variation and the speed at which drug resistance is acquired during treatment. We compared MIC values within all serial and recurrent cases from individual patients to determine how often MIC values differ between related isolates. Two-fold differences in MIC values (e.g., a single dilution) are not considered significant by CLSI or EUCAST standards due to inter-laboratory variation^54^. We identified nine serial isolate cases with a 4-fold to 64-fold variation in MIC (Supplementary Table S4). Notably, two of these cases had MIC differences to multiple drugs. Patient 54, a serial case of 8 *C. glabrata* isolates, had a 4-fold increase in amphotericin B MIC and a 64-fold increase in fluconazole MIC across the isolates, indicating substantial within-host variation of resistance (Figure 6A). Case 34, a *C. lusitaniae* serial case of 20 isolates, had an 8-fold increase in amphotericin B MIC and a 64-fold increase in micafungin MIC (Figure 6B). We also identified a 4-fold increase in amphotericin B MICs in two *C. albicans* cases, and a 4-fold to 8-fold increase in fluconazole MIC in three *C. glabrata* cases, one *C. utilis* case and one *C. parapsilosis* case. Differences in MIC values in serial isolates might represent existing within-host diversity of a strain or might be changes that are actively being selected for during antifungal therapy.

**Figure 6.**
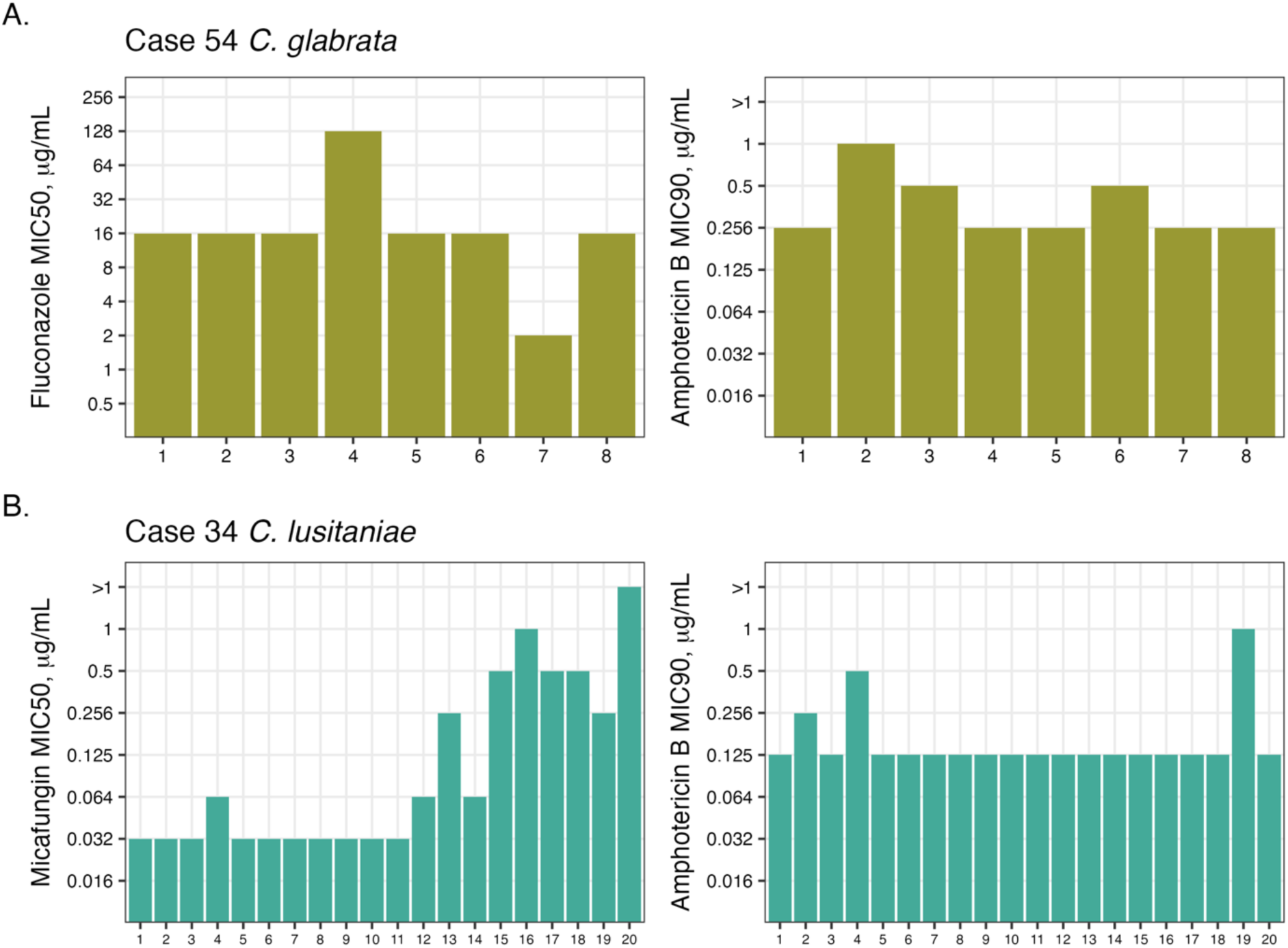
Within-host variation of MIC values occurs in all three drugs. (A). 64-fold fluconazole MIC differences and 4-fold amphotericin B MIC differences in case 54 *C. glabrata* isolates. (B.) 64-fold micafungin MIC differences and 8-fold amphotericin B in case 34 *C. lusitaniae* isolates.

### Fluconazole tolerance is greatest in *C. glabrata*

To evaluate antifungal tolerance, we determined 48-hour SMG values in fluconazole (Figure 7). SMG is calculated as the average growth across supra-MIC concentrations, relative to a no-drug control, and in *C. albicans* SMG values > 0.3 has been associated with persistent infections^35^. The *C. albicans* isolates in our study had a mean SMG of 0.2, with a range from 0.09 to 0.38. There were two recurrent *C. albicans* cases and all isolates had mean SMG values below 0.2, suggesting that fluconazole tolerance was unlikely to play a role in the recurrence.

**Figure 7.**
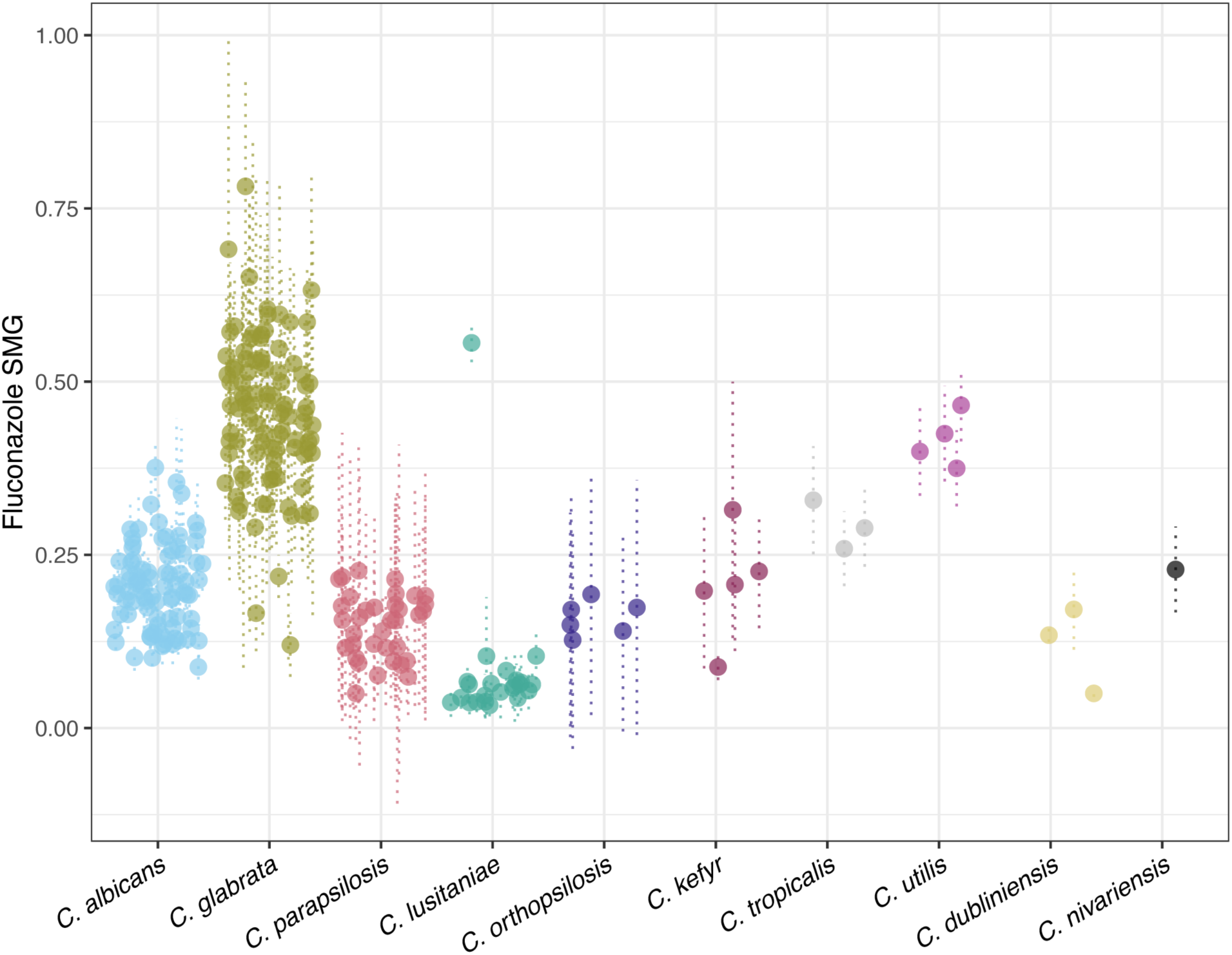
Fluconazole tolerance varies across and within *Candida* species. Dotplot of supra-MIC growth (SMG). For each isolate, the mean SMG is represented as a point and standard deviation is shown as dotted lines. SMG is the proportion of growth at 48 hours in all drug concentrations above the MIC, relative to a no-drug control. SMG testing was performed in triplicate for all isolates.

*C. glabrata* isolates had the highest fluconazole tolerance of all species in our study, with a mean SMG of 0.497 and range from 0.16 to 0.78 (Table S5). Despite the high SMG values, there were no recurrent *C. glabrata* cases in our study, and 19 of 20 serial isolate cases had time spans of less than 5 days. The *C. glabrata* isolates had high levels of fluconazole resistance along with tolerance, however their MIC and SMG values were not correlated (Spearman’s rank correlation coefficient = 0.12, p = 0.309 after multiple test correction), indicating that these are independent mechanisms of growth in the presence of drug.

*C. parapsilosis* isolates had generally low SMG values, with a mean of 0.15 and range from 0.05 to 0.23, which may indicate low tolerance but could also reflect overall slower growth in this species. Among the rare species, *C. lusitaniae* had the lowest fluconazole tolerance overall with a mean SMG of 0.08, but a single isolate had an SMG of 0.56, exceeding the tolerance of all species other than *C. glabrata*. Notably, this fluconazole tolerant *C. lusitaniae* isolate was also resistant to micafungin and amphotericin B. When comparing SMG values of isolates within serial cases, we identified multiple instances of SMG differences ≥ 0.1 involving *C. glabrata*, *C. albicans*, *C. parapsilosis*, *C. lusitaniae* and *C. kefyr* (Supplementary Table S6). These within-patient differences in tolerance may reflect existing phenotypic variation in a strain or may be evidence of within-host evolution during treatment – our data again highlights the value of testing multiple isolates from a patient over several days during antifungal treatment.

### There is limited association between growth rates in the absence of drug and MIC values

Bacteria often have a fitness cost associated with antimicrobial resistance, but in fungi the relationship between fitness and antifungal drug resistance is not straightforward^55^. We calculated the growth rate (r) of all isolates as a proxy for fitness over 24-hours in the absence of drug (Figure 8). *C. glabrata* and *C. nivariensis* had the fastest overall growth rates (*C. glabrata* median r = 0.988, *C. nivariensis* r = 1.16). *C. parapsilosis* had the slowest growth of any *Candida* species (median r = 0.373).

**Figure 8.**
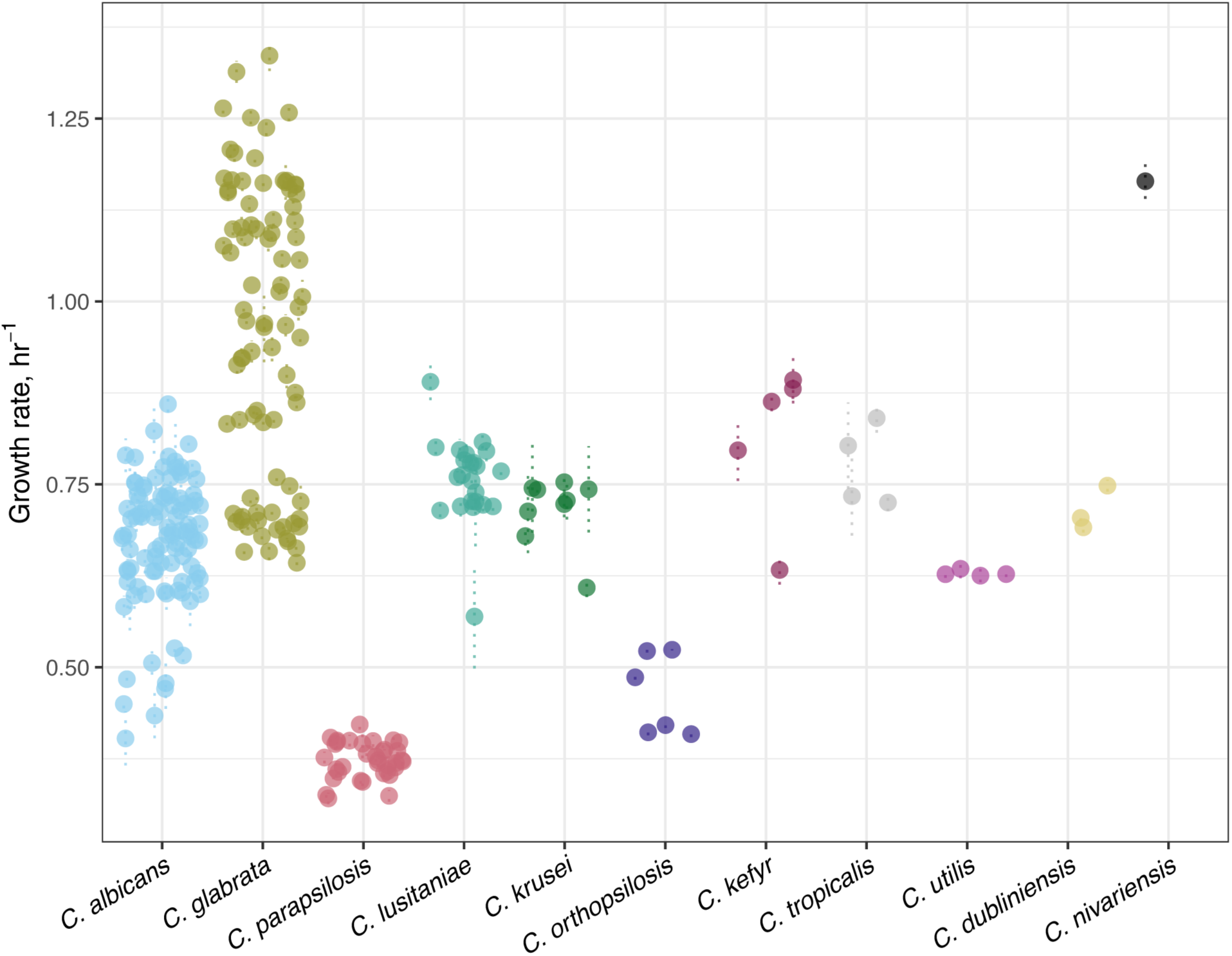
Distribution of isolate growth rate/hr^-1^ in the absence of drug. For each isolate, the mean growth rate in YPAD is represented as a point and standard deviation is shown as dotted lines. Growth curves were performed in triplicate for all isolates.

To determine if increased MIC values are associated with a growth defect, we tested whether there was a correlation between the MIC (in each drug class) and mean growth rate (in the absence of drug) for individual species. Species with a single MIC value for a given drug were excluded. In fluconazole, we identified a significant negative correlation between growth rate and MIC in *C. glabrata* (Spearman’s rank correlation coefficient = -0.24, p = 0.024) and in *C. utilis* (Spearman’s rank correlation coefficient = -1, p = 0), but no correlation in *C. albicans*, *C. parapsilosis*, and *C. tropicalis* (Supplementary Figure S1, Supplementary Table S7). In micafungin, there was no significant correlation between growth rate and MIC for all tested (Supplementary Figure S2, Supplementary Table S7). In amphotericin B, we identified a significant negative correlation between growth rate and MIC value in *C. orthopsilosis* (Spearman’s rank correlation coefficient = -0.87, p = 0.021). Surprisingly, in amphotericin B we identified a significant positive correlation between growth rate in the absence of drug and MIC for *C. glabrata* (Spearman’s rank correlation coefficient = 0.22, p = 0.039) and *C. parapsilosis* (Spearman’s rank correlation coefficient = 0.49, p = 0.003) (Supplementary Figure S3, Supplementary Table S7). There was no correlation in *C. albicans*, *C. lusitaniae*, *C. kefyr*, *C. utilis* or *C. dubliniensis*. Overall, our results indicate that the association between MIC and growth defects varies by species and by drug class, and that reduced drug susceptibility does not always confer a fitness cost.

## DISCUSSION

Candidemia is an important hospital-associated infection with significant associated mortality. Population-based surveillance studies have revealed important geographic and temporal variation in causal species and frequencies of antifungal drug resistance. We present the first study reporting species distribution and antifungal resistance for an academic medical center and 5 affiliated hospitals in the Twin Cities metro area. We prospectively collected 288 bloodstream isolates representing 11 *Candida* species from 119 patients, including 57 serial isolate cases representing 9 species.

*C. albicans*, *C. glabrata*, and *C. parapsilosis* were the most frequently recovered species in our study. Our data is relatively consistent with nation-wide studies from 2012 – 2017, albeit with slightly lower frequencies of C. *parapsilosis* and *C. tropicalis*^9,56^. Approximately 10% of our cases were caused by rare *Candida* species, highlighting their growing clinical importance. The frequency of polyfungal infections in our study (3.4%) is consistent with results reported by other studies in the United States and elsewhere^9,57^. The frequency of recurrent infection in this study, 3.4%, is somewhat lower than the 6% recurrence reported by a CDC EIP candidemia study of Georgia, Maryland, Oregon and Tennessee, which could reflect regional and temporal variation^58^.

We determined the frequency of antifungal drug resistance for all isolates to three drug classes. We identified no drug resistance in *C. albicans*, which is consistent with very low levels reported by the CDC EIP at four national sites^9^. We also identified no resistance to any drug classes in *C. parapsilosis*. Other U.S. studies have reported higher rates of fluconazole resistance in *C. parapsilosis* bloodstream isolates^9,59^. Local fluconazole resistance might be lower due to differences in treatment practices or might reflect undersampling due to the limited number of *C. parapsilosis* cases in this study^60^. We identified amphotericin B resistance in only a single *C. lusitaniae* isolate, which is consistent with low levels of amphotericin B resistance reported across *Candida* species^18,54^. Notably, this occurred within six days of initiation of therapy, highlighting how rapidly *C. lusitaniae* can acquire resistance to amphotericin B^61,62^.

Importantly, in *C. glabrata*, we identified fluconazole resistance in 16.7% of cases, micafungin resistance in 7.1% of cases and multidrug resistance in 4.8% of cases – frequencies that are 1.5 – 2 times higher than earlier studies of *C. glabrata* in the United States^9,45^. These results are clinically important and concerning because echinocandins are recommended first-line therapy for candidemia and fluconazole is an important step-down treatment for candidemia^63^. Our results may represent geographic variation or could reflect changes in treatment practices – in 2016, treatment guidelines for candidemia were revised to recommend echinocandins as first-line therapy, in contrast to the 2009 guidelines which indicated fluconazole was acceptable first-line treatment^63,64^. We identified multidrug resistance in a single case of *C. lusitaniae,* and we identified fluconazole resistance in one *C. utilis* case. Our results are an important addition to the very limited data available for both species. Overall, our findings emphasize that continued region-specific monitoring of antifungal drug resistance is crucial for identifying trends in resistance patterns that could impact antifungal stewardship efforts.

A strength of our study is the collection and testing of serial isolates from individual patients. We identified greater than 2-fold MIC differences in 9 of the 56 patients that had multiple isolates collected. In some cases, isolates collected on the same day had differing MIC values, while in other cases, MIC values climbed later during the infection or even dropped relative to earlier isolates. Our data highlight the potential clinical significance of within-host diversity and the limitations of current clinical testing strategies.

Tolerance has been associated with persistent or recurrent infection and is understudied in all *Candida* species^34^. We measured SMG as an indicator of fluconazole tolerance for all isolates and have reported some of the first SMG values for non-*albicans* species. While other studies have identified increased SMG values in persistent *C. albicans* infections, we did not find an association with SMG and recurrence^35^. However, our study collected a limited number of recurrent infections. Further investigation of tolerance mechanisms, including further longitudinal sampling and testing of additional drugs is an important direction for future work.

We determined the growth rate in rich media for all species as a measure of fitness in the absence of drug. There was extensive variation between isolates in *C. albicans* and *C. glabrata*, the two most common species in our study. Among the species that had a range of MIC values, we identified both positive and negative associations between increases in MIC and potential growth defects in the absence of drug. Our results suggest that there is not always a fitness trade-off associated with drug resistance in *Candida* species and highlight the need for further investigation to elucidate the relationship between acquired resistance and fitness in clinically relevant environments.

There are several limitations to our study. While serial isolates are well represented in our data set, all isolates are single colony subcultures. Our sampling strategy, while reflective of modern clinical microbiology practices, may underestimate the diversity of bloodstream populations during infection. Unstable genomic alterations such as aneuploidy, which can be important drivers of drug resistance and tolerance^38,65–67^, may be missed as a result of this single colony subculturing.

In conclusion, we surveyed species distribution and antifungal susceptibility of *Candida* bloodstream isolates in an academic health center and 5 affiliated hospitals. We identified important and deeply concerning trends in antifungal drug resistance in *C. glabrata* with implications for antifungal stewardship efforts. We have provided valuable phenotypic data for rare *Candida* species and described within-host phenotypic variability among common and rare *Candida* pathogens, which has potential clinical significance and is an important avenue for future research.

## METHODS

### IRB review

The study was reviewed and approved by the University of Minnesota Institutional Review Board (IRB ID STUDY00006473).

### Isolate and data collection

All available *Candida* bloodstream isolates identified from patients in M Health Fairview System hospitals were collected between December 2019 and May 2021. The M Health Fairview Infectious Diseases Diagnostic Laboratory performed species-level identification of all isolates by matrix-assisted laser desorption/ionization time-of-flight (MALDI-TOF). Each isolate collected for this study is a single colony subculture of an individual clinical blood culture. Isolates and patients were assigned unique study codes unrelated to identifying information. Colonies were cultured on Sabouraud dextrose agar plates and stocks were prepared with 20% glycerol and stored at -80°C.

### Minimum Inhibitory Concentration (MIC) by broth microdilution

MIC was determined by broth microdilution performed in RPMI 1640 (Cytiva, product no. SH30011.02) with 0.2% dextrose buffered with 0.165 M MOPS (Thermo Fisher Scientific, product no. J19256A1) and adjusted to pH 7.0^33^. Cultures were grown from glycerol stocks on Sabouraud dextrose agar plates at 35°C for 48 hours. Inoculum was prepared by suspending multiple colonies into sterile water and diluted to a final OD600 of 0.01. 100 µl of the inoculum was added to 96-well plates containing 100 µl of twofold serial dilution of antifungal drug in 2X RPMI medium (fluconazole: Alfa Aesar product no. J62015, micafungin: MedChemExpress product no. HY-17579, amphotericin B: Chem-Impex International product no. 00329). Plates were incubated in a humidified chamber at 35°C without shaking. At 24 hours post-inoculation, plate cultures were resuspended and OD530 readings were performed using a BioTek Epoch2 plate reader (Agilent). The mean and standard deviation of all 24-hour no-drug control OD530 readings were calculated per isolate from all plates. The EUCAST Antifungal MIC Methods for Yeast defines the MIC for azoles and echinocandins as the lowest drug concentration that inhibits ≥50% of growth relative to no-drug control, and the MIC for amphotericin B as the lowest concentration that inhibits ≥90% of growth relative to no-drug contro^33^. MICs for isolates with a no-drug control OD530 of > 0.2 were determined according to EUCAST guidelines and interpreted according to available EUCAST breakpoint values. Per EUCAST guidelines, isolates with an OD530 ≤ 0.2 were re-incubated per EUCAST guidelines and re-read at 48 hours. Isolates with an OD530 ≤ 0.2 at 48 hours were re-tested by gradient diffusion strip (see below). Quality control for each MIC batch was performed using *C. lusitaniae* FDA-CDC AR Bank # 0398 and/or *C. krusei* FDA-CDC AR Bank # 0397^68^. All MIC assays were performed in triplicate.

### MIC by gradient diffusion strip

MIC testing by gradient diffusion strip was performed for isolates with no-drug control OD values ≤ 0.2 at 24 and 48 hours. Antifungal susceptibility testing was adapted from CLSI supplement M60 document protocol for Gradient Diffusion Strips^69^. Briefly, isolates were struck from glycerol stocks onto Sabouraud dextrose agar plates and incubated for 24 hours at 35°C. For each culture, multiple colonies were picked, suspended in sterile water and diluted to a final OD600 of 0.01 using a spectrophotometer. 100µl of the diluted cells was plated onto RPMI plates, a gradient diffusion strip (fluconazole: Biomerieux, product no. 510858, amphotericin B: Liofilchem, product no. 921531) was applied and the plate was incubated for 24 hours at 35°C in a humidified chamber. At 24 hours, plates were imaged using a Bio-Rad Gel Doc system. The MIC values were determined by identifying the concentration where the lawn of growth intersected with the gradient strips.

### Supra-MIC growth (SMG) by broth microdilution

SMG was calculated for all isolates with no-drug control OD values > 0.2 at 24 hours. Plates were incubated for an additional 24 hours at 35°C. Plate cultures were resuspended and 48-hour OD530 readings were performed using a BioTek Epoch2 plate reader (Agilent). Supra-MIC growth (SMG) in fluconazole was calculated as the mean of 48-hour growth in all wells above the MIC concentration, divided by the mean of the no-drug control wells. All SMG assays were performed in triplicate.

### Growth curve analysis

Overnight cultures were started from glycerol stocks and grown in a shaking incubator at 30°C in liquid YPAD medium with 2% dextrose (10 g/L yeast extract, 20 g/L Bacto peptone, 20 g/L dextrose, 0.04 g/L adenine and 0.08 g/L uridine). Overnight cultures were diluted in fresh YPAD medium to a final OD600 of 0.01 and 20 µl of this cell suspension was inoculated into a 96-well plate containing 180 µl of YPAD with 1% dextrose. Cells were grown in a BioTek Epoch2 plate reader at 30°C for 24 hours with constant shaking, and OD600 readings were taken every 15 minutes. Growth curves were performed in triplicate. Growth curve metrics including mean and standard deviation for carrying capacity, growth rate, doubling time, AUC-E and AUC-L were calculated with the R package *Growthcurver* (v0.3.1)^70,71^. Metrics were plotted with the R package *ggplot2* (v3.5.1)^72^.

### Correlation testing

For species that had more than one MIC value, Spearman’s rank correlation coefficient was calculated for MIC relative to SMG and to growth rate in YPAD. All correlation analyses and multiple test correction (using the Holm method) were performed with the R package *correlation* (v0.8.4)^73,74^.

### Code availability

Scripts used in data analysis and figure generation are available at https://github.com/selmeckilab/2024_Candida_clinical_isolate_phenotyping.

## ACKNOWLEDGEMENTS

We are grateful to members of the Selmecki laboratory for feedback on data analysis, figures and the manuscript. This work was supported by the National Institutes of Health (R01AI143689), National Science Foundation (DBI-232051), and Burroughs Wellcome Fund Investigator in the Pathogenesis of Infectious Diseases Award (#1020388) to A.S., the Undergraduate Research Opportunity (UROP) to C.Z., the University of Minnesota Graduate School Doctoral Dissertation Fellowship (DDF) to N.E.S. The Minnesota Supercomputing Institute (MSI) at the University of Minnesota provided resources that contributed to the research results reported within this paper.

